# Cell type specific genetic regulation of gene expression across human tissues

**DOI:** 10.1101/806117

**Authors:** Sarah Kim-Hellmuth, François Aguet, Meritxell Oliva, Manuel Muñoz-Aguirre, Valentin Wucher, Silva Kasela, Stephane E. Castel, Andrew R. Hamel, Ana Viñuela, Amy L. Roberts, Serghei Mangul, Xiaoquan Wen, Gao Wang, Alvaro N. Barbeira, Diego Garrido-Martín, Brian Nadel, Yuxin Zou, Rodrigo Bonazzola, Jie Quan, Andrew Brown, Angel Martinez-Perez, José Manuel Soria, GTEx Consortium, Gad Getz, Emmanouil T. Dermitzakis, Kerrin S. Small, Matthew Stephens, Hualin S. Xi, Hae Kyung Im, Roderic Guigó, Ayellet V. Segrè, Barbara E. Stranger, Kristin G. Ardlie, Tuuli Lappalainen

## Abstract

The Genotype-Tissue Expression (GTEx) project has identified expression and splicing quantitative trait loci (*cis*-QTLs) for the majority of genes across a wide range of human tissues. However, the interpretation of these QTLs has been limited by the heterogeneous cellular composition of GTEx tissue samples. Here, we map interactions between computational estimates of cell type abundance and genotype to identify cell type interaction QTLs for seven cell types and show that cell type interaction eQTLs provide finer resolution to tissue specificity than bulk tissue *cis*-eQTLs. Analyses of genetic associations to 87 complex traits show a contribution from cell type interaction QTLs and enables the discovery of hundreds of previously unidentified colocalized loci that are masked in bulk tissue.

**One Sentence Summary:** Estimated cell type abundances from bulk RNA-seq across tissues reveal the cellular specificity of quantitative trait loci.

## Main Text

The Genotype-Tissue Expression (GTEx) project (*1*) and other studies (*2-5*) have shown that genetic regulation of the transcriptome is widespread. GTEx in particular has built an extensive catalog of expression and splicing quantitative trait loci in *cis* (*cis*-eQTLs and *cis*-sQTLs) across an unprecedented range of tissues, showing that QTLs are generally either highly tissue-specific or widely shared, even across dissimilar tissues and organs (*1, 6*). However, the vast majority of these studies have been performed using heterogeneous bulk tissue samples comprising diverse cell types. This limits the power, interpretation, and downstream applications of QTL studies. Genetic effects that are active only in rare cell types may be left undetected, mechanistic interpretation of QTL sharing across tissues and other contexts is complicated without understanding differences in cell type composition, and inference of downstream molecular effects of regulatory variants without the specific cell type context is challenging. Efforts to map eQTLs in individual cell types have been largely restricted to blood, using purified cell types (*7-10*) or single cell sequencing (*11*). Cell type specific eQTLs can also be computationally inferred from bulk tissue measurements, using the estimated proportion or enrichment of relevant cell types to test for an interaction with genotype, but such approaches to date have been applied to only a limited range of cell types, including blood cell types (*12, 13*) and adipocytes (*14*). These studies identified thousands of cell type interactions in eQTLs discovered in whole blood samples from large cohorts [5,683 samples (*12*); 2,116 samples, (*13*)], indicating that large numbers of interactions are likely to be identified by expanding this type of analysis to other tissues and cell types.

In this study, we applied cell type deconvolution to characterize the cell type specificity of *cis*-eQTLs and *cis*-sQTLs for 43 cell type-tissue combinations, using seven cell types across 35 tissues (Fig. 1A). Estimating the cell type composition of a tissue biospecimen from RNA-seq remains a challenging problem (*15*) and multiple approaches for inferring cell type proportions have been proposed (*16*). We performed extensive benchmarking for multiple cell types across several expression datasets (fig. S1). The xCell method (*17*), which estimates the enrichment of 64 cell types using reference profiles, was most robust based on correlation with cell counts in blood (fig. S1A), *in silico* simulations (fig. S1B), and correlation with expression of marker genes for each cell type (fig. S1C). Furthermore, the inferred abundances reflected differences in histology and tissue pathologies (fig. S1D, E). For each cell type, we selected tissues where the cell type was highly enriched to map cell type interacting eQTLs in *cis* (fig. S2A, B). The xCell scores for these tissue-cell type pairs were also highly correlated with the PEER factors used to correct for unobserved confounders in the expression data for QTL mapping (*1*) (fig. S2C), suggesting that cell type composition likely explains a large part of inter-sample variation in gene expression.

**Fig. 1.**
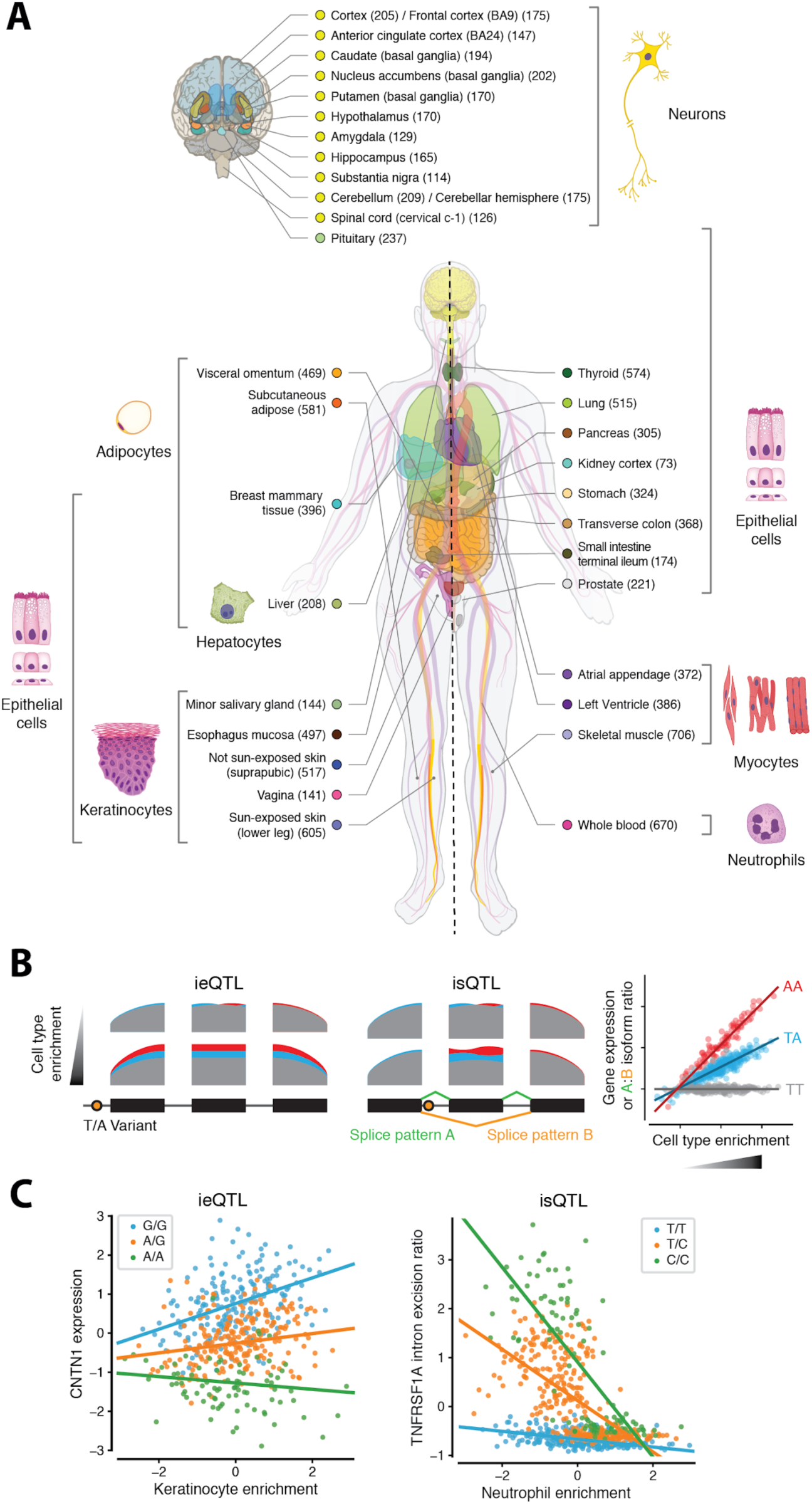
Study design of mapping cell type ieQTLs and isQTLs in GTEx v8 project. (**A**) Illustration of 43 cell type-tissue pairs included in the GTEx v8 project. Cell types with median xCell enrichment score > 0.1 within a tissue were used (fig. S2). (**B**) Schematic representation of a cell type interacting eQTL and sQTL. (**C**) Example cell type ieQTL and isQTL. The *CNTN1* eQTL effect in not sun-exposed skin is associated with keratinocyte abundance (left panel). The *TNFRSF1A* sQTL effect in whole blood is associated with neutrophil abundance, but is only detected in samples with lower neutrophil abundances (right panel). Each data point represents an RNA-seq sample and is colored by the ieQTL and isQTL genotypes, respectively. The regression lines correspond to the coefficients of the interaction model.

We used a linear regression model for gene expression that included an interaction between cell type enrichment and genotype, thus using variability in cell type composition between individuals to identify eQTLs whose effect varies depending on the enrichment of the cell type (Fig. 1B). Since QTLs identified this way are not necessarily specific to the estimated cell type but may reflect another (anti)correlated cell type, we refer to these eQTLs as cell type interacting eQTLs, or cell type ieQTLs. We applied an analogous approach to map cell type interacting splicing QTLs (isQTLs), using intron excision ratios that reflect alternative isoform usage, quantified by LeafCutter (*18*) (Fig. 1B). Across cell types and tissues, we detected 3,347 protein coding and lincRNA genes with an ieQTL (ieGenes) and 987 genes with an isQTL (isGenes) at 5% FDR per cell type-tissue combination (Fig. 2A, fig. S3A+B and table S1). In the following analyses, ieQTLs and isQTLs with 5% FDR are used unless indicated otherwise. The QTL effect of ieQTLs and isQTLs can increase or decrease as a function of cell type enrichment (Fig. 1C, fig. S3C+D). This correlation is usually positive (56%; median across cell type-tissue combinations); for example, a keratinocyte ieQTL for *CNTN1* in skin had a particularly strong effect in samples with high enrichment of keratinocytes. However, for a significant number of ieQTLs the effect was negatively correlated (18%) or ambiguous (24%) (fig. S4A,B), with the interaction likely capturing a QTL that is active in another cell type. Notably, while 85% of ieQTLs corresponded to genes with at least one standard eQTL, 21% of these ieQTLs were not in LD (R^2^ < 0.2) with any of the corresponding eGene’s conditionally independent eQTLs (fig. S4C), indicating that ieQTL analysis often reveals genetic regulatory effects that are not detected by standard eQTL analysis of heterogeneous tissue. Unlike for bulk tissue *cis*-QTLs, iQTL discovery was only modestly correlated with sample size (Spearman’s ρ = 0.53 and 0.35, respectively; fig. S3E+F). The tissues with most iQTLs included blood, as well as breast and transverse colon that both stratified into at least two distinct groups based on histology (*19*): epithelial vs. adipose tissue (breast) and mucosal vs. muscular tissue (colon) (fig. S1B). This suggests that high inter-individual variance in cell type enrichments driven by tissue heterogeneity is a major determinant in discovery power and benefits iQTL mapping despite being a complicating factor for many other types of tissue gene expression analyses. Downsampling analyses in whole blood and transverse colon revealed linear relationships between sample size and ieQTL discovery in these tissues, suggesting that significantly larger numbers of ieQTLs may be discovered with larger sample sizes (fig. S3G).

**Fig. 2.**
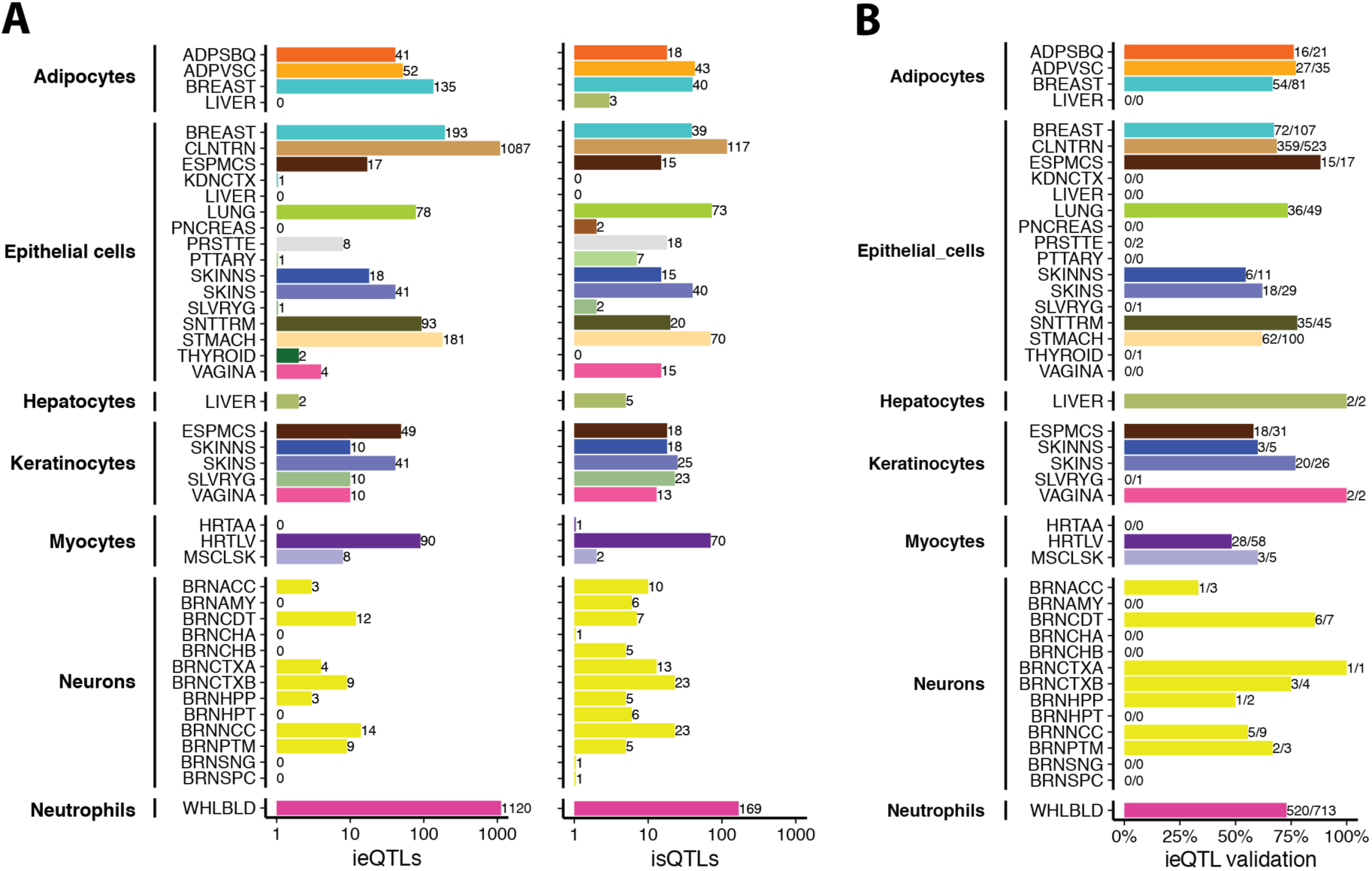
Cell type ieQTL and isQTL discovery. (**A**) Number of cell type ieQTLs (left panel) and isQTLs (right panel) discovered in each cell type-tissue combination at FDR < 5%. Bar labels show the number of ieQTLs and isQTLs, respectively. See Fig. 1A for the legend of tissue colors. (**B**) Proportion of cell type ieQTLs that validated in ASE data. Validation was defined as ieQTLs for which the Spearman correlation between allelic fold-change (aFC) estimates from ASE and cell type estimates was nominally significant (*P* < 0.05). Tissue abbreviations are provided in table S2. Bar labels indicate the number of ieQTLs with validation/number of ieQTLs tested.

Since external replication data sets are sparse, we used allele-specific expression (ASE) data of eQTL heterozygotes (*20, 21*) to correlate individual-level quantifications of the eQTL effect size (measured as allelic fold-change, aFC) with individual-level cell type enrichments. If the eQTL is active in the cell type of interest, we expect to see low aFC in individuals with low cell type abundance, and higher aFC in individuals with high cell type abundance (fig. S5A). Spearman correlation p-values can then be used to assess how many cell type ieQTLs show evidence of validation using this approach. The median proportion of ieQTLs with a significant aFC-cell type correlation (*P* < 0.05) was 0.63 (Fig. 2B). For 13 cell type-tissue combinations with > 20 significant ieQTLs, the π_1_ statistic corresponding to the correlation p-values (*22*) confirmed the high validation rate (mean π_1_ = 0.76, fig. S5B). While this approach does not constitute formal replication in an independent cohort, it is applicable to all tested cell type-tissue combinations, and corroborates that ieQTLs are not statistical artefacts of the interaction model. Next, we performed replication analyses in external cohorts, including whole blood from the GAIT2 study (*23*), purified neutrophils (*8*), adipose and skin tissues from the TwinsUK study for ieQTLs (*5*) and temporal cortex from the Mayo RNA-sequencing study for both ieQTLs and isQTLs (*24*). Overall replication was moderate to high (π_1_ = 0.32 - 0.67) with the highest replication rates observed in purified neutrophils for whole blood (fig. S6A+E). The differences in replication rates likely reflect a combination of lower power to detect cell type ieQTLs/isQTLs compared to standard eQTLs/sQTLs, as well as differences in tissue heterogeneity across studies. Taken together, these results show that ieQTLs and isQTLs can be detected with reasonable robustness for diverse cell types and tissues.

Next, we sought to determine to what extent cell type ieQTLs contribute to the tissue specificity of *cis*-eQTLs. First, we analyzed ieQTL sharing across cell types, observing that ieQTLs for one cell type were generally not ieQTLs for other cell types (e.g., myocyte ieQTLs in muscle tissues were not hepatocyte ieQTLs in liver, etc.; fig. S7B). To determine if a significant cell type interaction effect is associated with the tissue-specificity of an eQTL, we tested whether cell type ieQTLs are predictors of tissue sharing. We annotated the top *cis*-eQTLs per gene across tissues with their cell type ieQTL status for the five cell types with at least 20 ieQTLs (adipocytes, epithelial cells, keratinocytes, myocytes, and neutrophils). This annotation was included as a predictor in a logistic regression model of eQTL tissue sharing based on eQTL properties including effect size, minor allele frequency, eGene expression correlation, genomic annotations, and chromatin state (*1*). In all five cell types, ieQTL status was a strong negative predictor of tissue-sharing, with the magnitude of the effect similar to that of enhancers, indicating that ieQTLs are an important mechanism for tissue-specific regulation of gene expression (Fig. 3A, fig. S7A). We corroborated this finding using multi-tissue eQTL mapping with MASH (*1*), testing whether eGenes that are tissue-specific (eQTLs discovered at LSFR < 0.05 only in the tissue/tissue type of interest) have a higher proportion of cell type ieQTLs compared to eGenes that are shared across tissues (LSFR < 0.05 in multiple tissues). Indeed, the proportion of cell type ieQTLs across all 43 cell type-tissue combinations was significantly higher in tissue-specific eGenes compared to tissue-shared eGenes (*P* = 1.9e-05, one-sided Wilcoxon rank sum test, Fig. 3B) further highlighting the contribution of cell type-specific genetic gene regulation to tissue specificity of eQTLs.

**Fig. 3.**
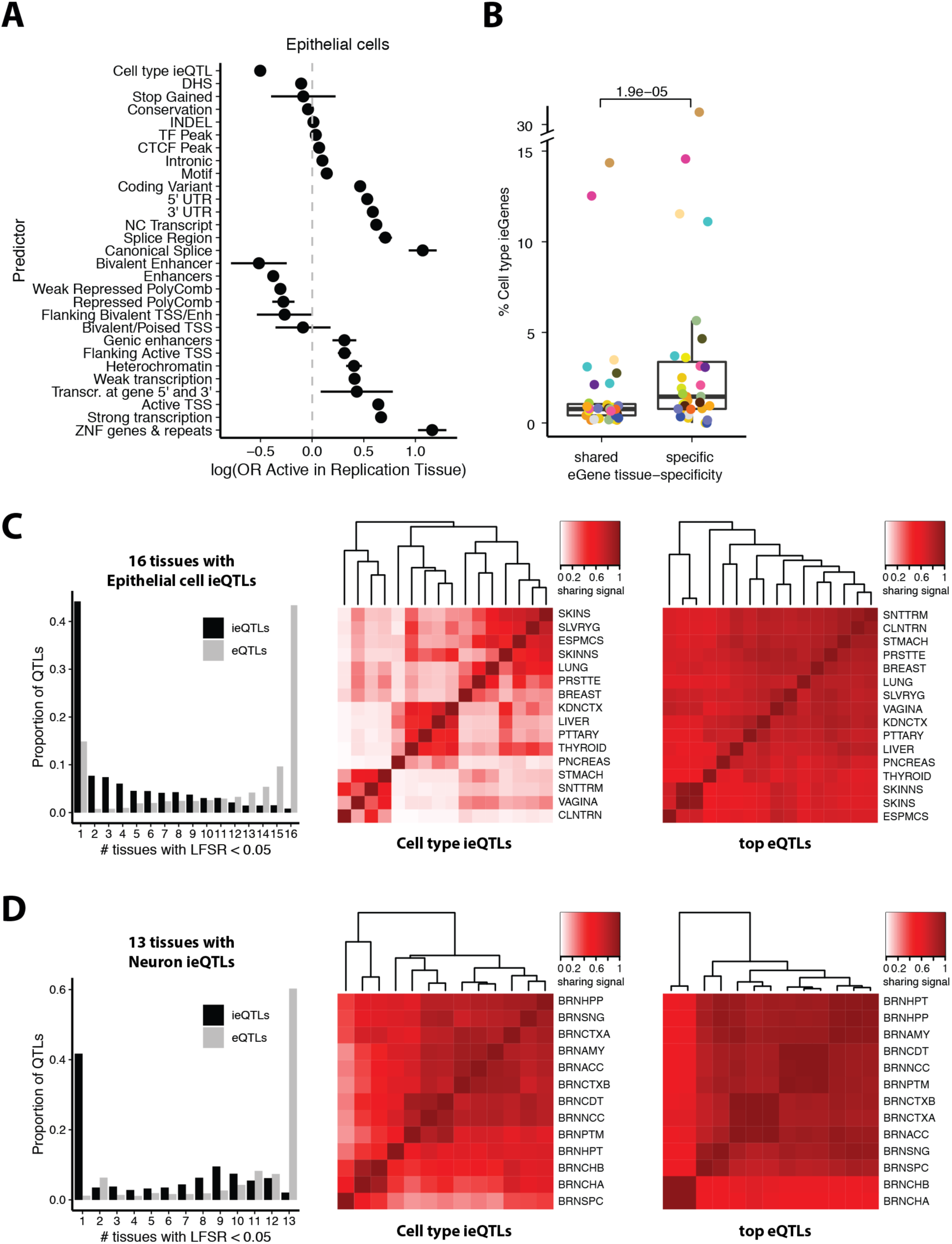
Cell type ieQTLs contribute to *cis-*eQTL tissue specificity. (**A**) Coefficients from logistic regression models of *cis-*eQTL tissue sharing, where epithelial cell ieQTL status is one of the predictors: All significant top *cis-* eQTLs per tissue were annotated based on if they were also a significant ieQTL for a given cell type. The coefficients represent the log(odds ratio) that an eQTL is active in a replication tissue given a predictor. Chromatin states were defined using matched Epigenomics Roadmap tissues and the 15-state ChromHMM (*25*). Genomic annotations, conservation, and overlaps with Ensembl regulatory build TF, CTCF, and DHS peaks are also included. Bars represent the 95% confidence interval. (**B**) Proportion of cell type ieQTL-genes (ieGenes) among tissue-specific and tissue-shared eGenes. An eGene is considered tissue-specific if its eQTL had a MASH local false sign rate (LFSR, equivalent to FDR) < 0.05 only in the cell type ieQTL tissue (or tissue type) otherwise it is considered tissue-shared. Results of all 43 cell type-tissue combinations are shown. See Fig. 1A for the legend of tissue colors. (**C+D**) Tissue activity of cell type ieQTLs and eQTLs, where a cell type ieQTL and eQTL was considered active in a tissue if it had an LFSR < 0.05 (left panel). Pairwise tissue-sharing of ieQTLs (middle panel) or lead standard *cis-*eQTLs (right panel) respectively. The color-coded sharing signal is the proportion of significant QTLs (LFSR < 0.05) that are shared in magnitude (within a factor of 2) and sign between two tissues.

To examine the sharing patterns of cell type ieQTLs across tissues we used two cell types with ieQTLs mapped in >10 tissues (16 tissues for epithelial cells and 13 for neurons). We observed that while standard eQTLs were highly shared across the subsets of 16 and 13 tissues, cell type ieQTLs tended to be highly tissue specific, reflected by an average of four and five tissues with shared ieQTL effects compared to 11 and 12 for eQTLs in epithelial and brain tissues respectively (Fig. 3C+D, left panels). 25.3% of neuron ieQTLs were shared between nine brain tissues, highlighting that tissues of the cerebrum (e.g., cortex, basal ganglia, limbic system) show particularly high levels of sharing compared to cerebellar tissues, the hypothalamus, and the spinal cord (Fig. 3D, left panel). This pattern was absent when analyzing standard eQTLs. Pairwise tissue sharing comparisons further confirmed that cell type ieQTLs showed greater tissue specificity and more diverse tissue sharing patterns than standard eQTLs, which were broadly shared across all tissues (Fig. 3C+D, middle and right panels). These results show that incorporating cell type composition is essential for characterizing the sharing of genetic regulatory effects across tissues.

To study the contribution of cell type interacting QTLs to GWAS associations of 87 complex traits, we first examined the enrichment of iQTLs of each cell type/tissue combination for trait associations (GWAS *P* ≤ 0.05) using QTLEnrich (v2) (*26*). We used 23 and 7 cell type/tissue pairs (19 and 7 unique tissues) with >100 ieQTLs or isQTLs at a relaxed FDR (40% FDR) to generate robust enrichment estimates of 87 GWAS traits. Across all tested cell type/tissue-trait pairs, the GWAS signal was clearly enriched among ieQTLs and isQTLs (1.3 and 1.4 median fold-enrichments, respectively), similarly to standard eQTLs and sQTLs (Fig. 4A, table S4). The GWAS enrichments were robust to the iQTL FDR cutoffs (Fig. S8A+B). We next analyzed the enrichments of the individual traits for iQTLs of two cell types with best power: neutrophil iQTLs in blood and epithelial cell iQTLs in transverse colon. We compared them to the corresponding standard QTLs (Fig. 4B, Fig S8C+D), focusing on traits that had a significant enrichment for either QTL type (Bonferroni-adjusted *P* < 0.05). Interestingly, in blood we observed a significant shift towards higher enrichment for ieQTLs (one-sided, paired Wilcoxon rank sum test; *P* = 0.0026) and especially isQTLs (*P* = 2.8e-05), which appears to be driven by GWAS for blood cell traits, and also immune traits having a higher enrichment for iQTLs. The higher iQTL signal is absent in colon (ieQTL *P* = 1 and isQTL *P* = 0.13), even though the standard QTL enrichment for blood cell traits appear similar for blood and colon. This pattern suggests that cell type interacting QTLs may have better resolution for indicating relevant tissues and cell types for complex traits, compared to tissue QTLs, but future studies are needed to fully test this hypothesis.

**Fig. 4.**
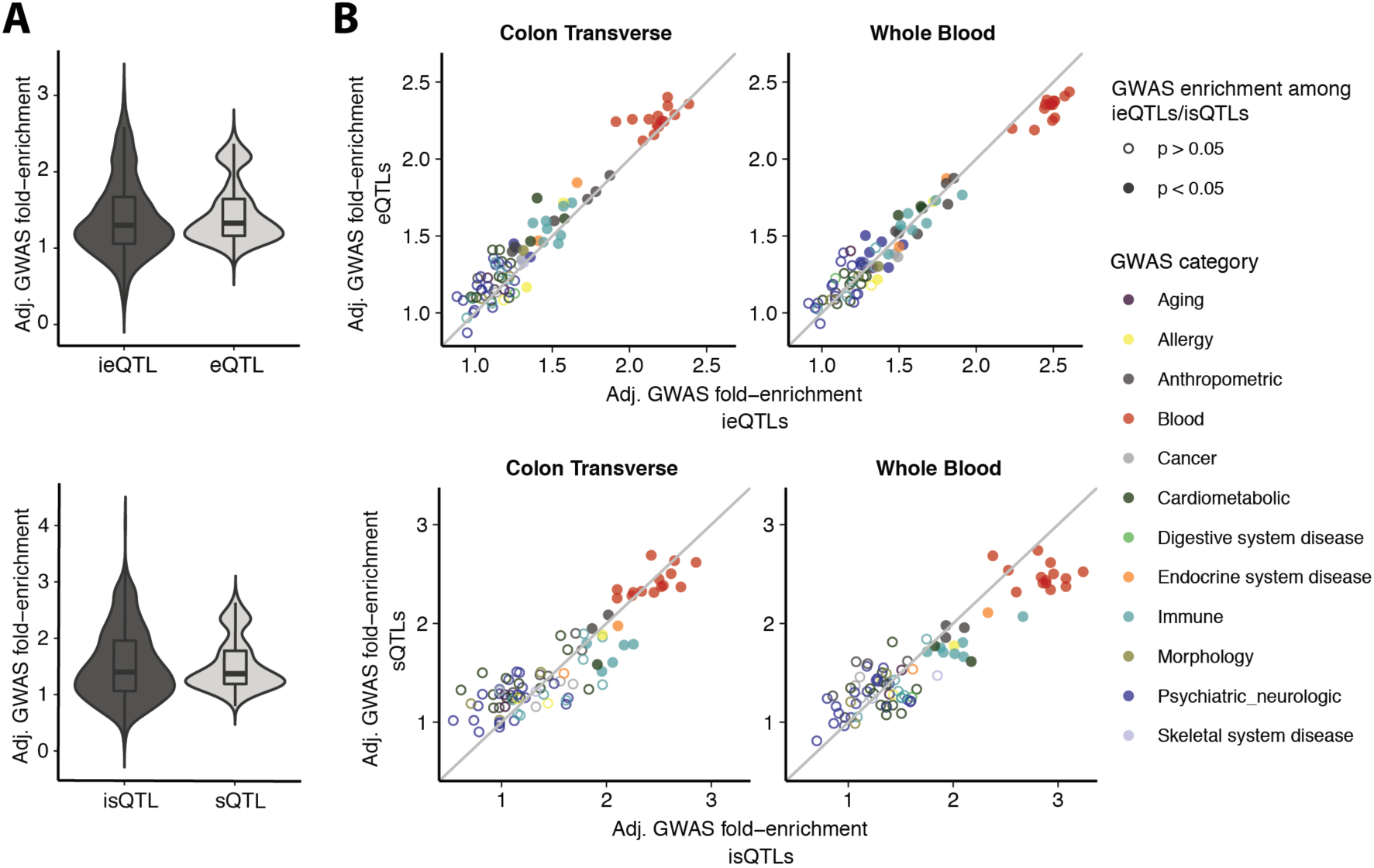
Cell type iQTLs are enriched for GWAS signals. **(A)** Distribution of adjusted GWAS fold-enrichment of 23×87 (top panel) and 7×87 (bottom panel) tissue-trait combinations using the most significant iQTL or standard QTL per eGene/sGene. (**B**) Adjusted GWAS fold-enrichments of 87 GWAS traits among iQTLs on the x-axis and standard QTLs on the y-axis. Filled circles indicate significant GWAS enrichment among iQTLs at *P* < 0.05 (Bonferroni-corrected). Colors represent GWAS categories of the 87 GWAS traits (see table S3).

We next asked whether cell type iQTLs can be linked to loci discovered in genome-wide association studies (GWAS) as well as pinpoint the cellular specificity of these associations. To this end, we tested 13,702 ieGenes and 2,938 isGenes (40% FDR) for colocalization with 87 GWAS traits (*1, 27*), using both the cell type ieQTL/isQTL and corresponding standard bulk tissue QTL. 1,370 (10.3%) cell type ieQTLs and 89 (3.7%) isQTLs colocalized with at least one GWAS trait (Fig. 5A, table S5+S6). The larger number of colocalizations identified for neutrophil ieQTLs and isQTLs in whole blood relative to other cell type-tissue pairs likely reflects a combination of the larger number of ieQTLs and isQTLs and the abundance of significant GWAS loci for blood-related traits in our set of 87 GWASs. Our analysis revealed a substantial proportion of loci for which only the ieQTL/isQTL colocalizes with the trait (467/1370, 34%), or where the joint colocalization of the ieQTL/isQTL and corresponding standard eQTL indicates the cellular specificity of the trait as well as its potential cellular origin (401/1370, 29%). For example, a colocalization between the *DHX58* gene in the left ventricle of the heart and an asthma GWAS was only identified through the corresponding myocyte ieQTL (PP4 = 0.64), but not the standard eQTL (PP4 = 0.00; Fig. 5B). Cardiac cells such as cardiomyocytes are not primarily viewed to have a causal role in asthma, but their presence along pulmonary veins and their potential contribution to allergic airway disease have been previously described (*28*). An example where both the standard eQTL and the cell type ieQTL colocalize with the trait is given in Fig. 5C for *KREMEN1* in adipocytes in subcutaneous adipose tissue and a birth weight GWAS (PP4 ∼0.8); *KREMEN1* has been linked to adipogenesis in mice (*29*). We highlight two analogous examples for isQTLs: the epithelial cell isQTL for *CDHR5* in small intestine colocalized with eosinophil counts whereas the standard sQTL did not (Fig. 5D), and conversely, both the standard sQTL and myocyte isQTL for *ATP5SL* in the left ventricle of the heart colocalized with standing height (Fig. 5E). While the iQTLs do not necessarily pinpoint the specific cell type where the regulatory effect is active, they indicate that cell type specificity plays a role in the GWAS locus. Together, the colocalization results show that cell type interaction QTLs yield new potential target genes for GWAS loci that are missed by tissue QTLs, and provide hypotheses of cellular specificity of regulatory effects underlying complex traits.

**Fig. 5.**
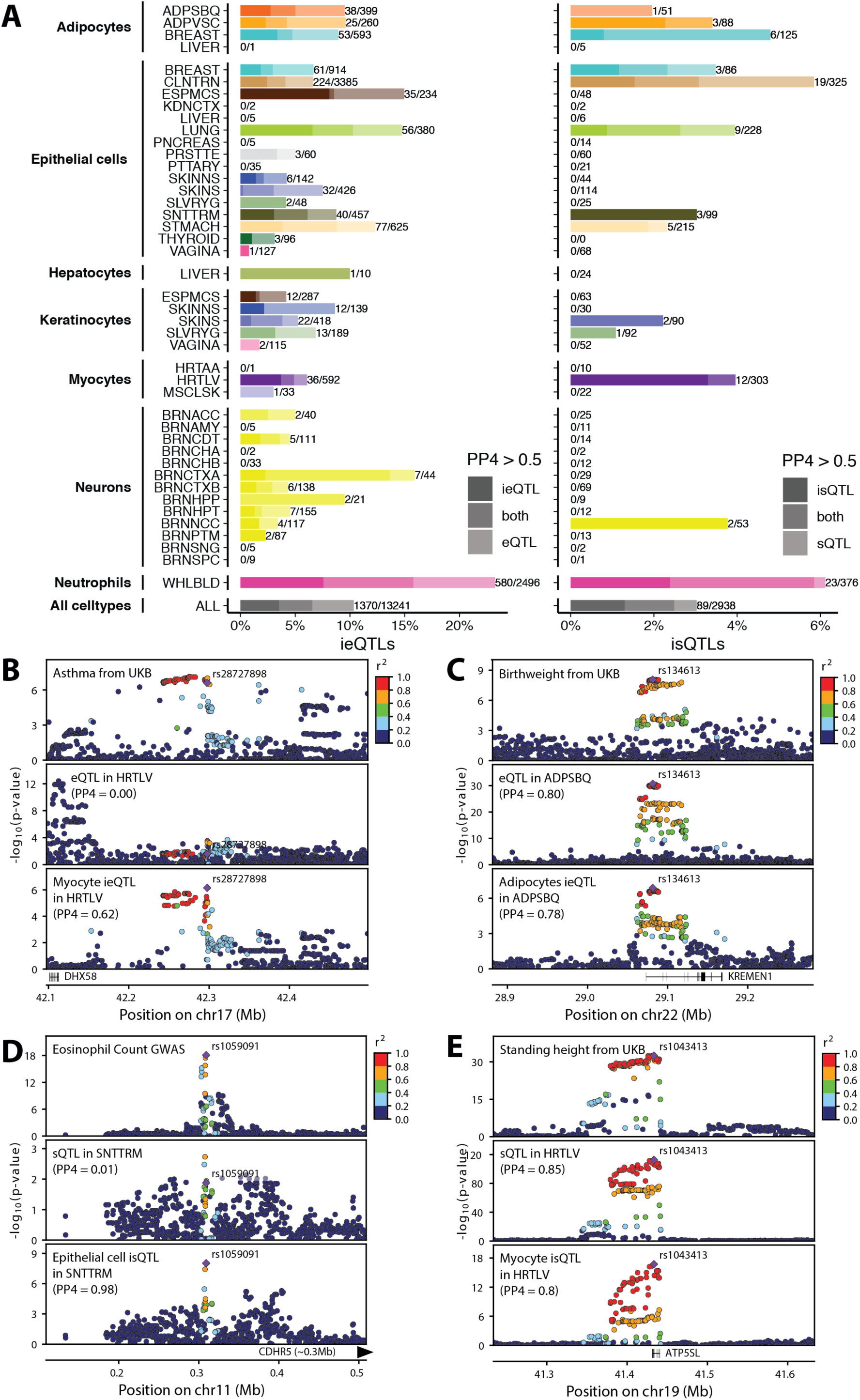
Cell type iQTLs improve GWAS-QTL matching. **(A)** Proportion of cell type ieQTLs (left panel) or isQTLs (right panel) with evidence of colocalization using COLOC posterior probabilities (PP4 > 0.5), for ieQTLs and isQTL at FDR < 0.4. Color saturation indicates if a trait colocalized with the cell type iQTL only (dark), the *cis*-QTL only (light) or both QTLs (medium). Bar labels indicate the number of cell type iQTLs with evidence of colocalization (either as iQTL or *cis*-QTL)/number of iQTLs tested. (**B**) Association p-values in the *DHX58* locus for an asthma GWAS (top panel), bulk heart left ventricle *cis*-eQTL (middle panel), and myocyte ieQTL (bottom panel). (**C**) Association p-values in the *KREMEN1* locus for a birth weight GWAS (top panel), bulk subcutaneous adipose *cis*-eQTL (middle panel), and adipocyte ieQTL (bottom panel). (**D**) Association p-values in the *CDHR5* locus for an eosinophil count GWAS (top panel), bulk small intestine terminal ileum *cis*-sQTL (middle panel), and epithelial cell isQTL (bottom panel). (**E**) Association p-values in the *ATP5SL* locus for a standing height GWAS (top panel), bulk heart left ventricle *cis*-sQTL (middle panel), and myocyte isQTL (bottom panel).

By mapping interaction effects between cell type enrichment and genotype on the transcriptome across GTEx tissues, we were able to identify thousands of eQTLs and sQTLs that are likely to be cell type specific. Notably, the ieQTLs and isQTLs we report here include immune and stromal cell types in tissues where cell type specific QTLs have not yet been characterized. Cell type ieQTLs are strongly enriched for tissue- and cellular specificity, and provide a finer resolution to tissue-specificity than bulk *cis*-eQTLs that are highly shared between tissues. It is likely that many more cell type ieQTLs remain to be discovered for cell types and tissues not considered in this study, and improving deconvolution approaches and sample sizes will be valuable in this effort. However, the substantial allelic heterogeneity observed in standard eQTLs (*1*) and limited power to deconvolve QTLs that are specific to rare cell types or with weak or opposing effects indicate that many more cell type specific eQTLs exist beyond those that can be computationally inferred from bulk tissue data. We therefore anticipate that single-cell QTL studies will be essential to complement the approaches presented here. Given the enrichment of GWAS signal in cell type iQTLs for cell types potentially relevant to the traits, and the large fraction of colocalizations with GWAS traits that are only found with cell type iQTLs, it will be essential to exhaustively characterize cell type specific QTLs to contribute towards a mechanistic understanding of these loci.

## Supporting information

Kim-Hellmuth et al supplement

## Acknowledgments

We thank the donors and their families for their generous gifts of organ donation for transplantation, and tissue donations for the GTEx research project; Mariya Khan and Christopher Stolte for the illustrations in Figure 1.

## Author contributions

S.K.-H., F.A. and T.L. conceived the study. S.K.-H. and F.A. led the writing, figure generation and editing of the manuscript and supplement. S.K.-H. coordinated analyses of all contributing authors; S.K.-H. and F.A. generated pipelines and performed iQTL mapping; S.K.-H., F.A., M.O., M.M.-A., V.W., D.G.-M., S.M., B.N., J.Q. performed cell type benchmarking analysis; S.K. performed ieQTL validation with ASE data using validation pipeline and ASE data generated by S.E.C; F.A., A.V., A.L.R. performed replication analysis; S.E.C. performed QTL tissue activity prediction analysis; S.K.-H. and S.E.C. generated tissue sharing (MASH) data; S.K.-H. performed tissue specificity, multi-tissue analysis and colocalization analysis; A.R.H. performed QTLEnrich analysis; G.W. and Y.Z. provided software support for multi-tissue eQTL analysis; X.W. and H.I. provided advice on colocalization analysis; A.B., A.M.-P., J.M.-S. contributed to replication analysis; F.A. and K.G.A. generated and oversaw GTEx v8 data generation, LDACC & pipelines; A.N.B. and R.B. generated GWAS data; K.S.S., M.S., H.S.X., G.G., E.T.D., H.I., R.G., A.V.S., B.E.S., K.G.A., T.L. supervised the work of trainees in his/her lab; M.O. and T.L. contributed to editing of the manuscript; All authors read and approved the final manuscript.

## Data and materials availability

All GTEx open-access data, including summary statistics and visualizations of cell type iQTLs, are available on the GTEx Portal (https://gtexportal.org/home/datasets). All GTEx protected data are available via dbGaP (accession phs000424.v8). Access to the raw sequence data is now provided through the AnVIL platform (https://gtexportal.org/home/protectedDataAccess). The QTL mapping pipeline is available at https://github.com/broadinstitute/gtex-pipeline, tensorQTL is available at https://github.com/broadinstitute/tensorqtl. Residual GTEx biospecimens have been banked, and remain available as a resource for further studies (access can be requested on the GTEx Portal, at https://www.gtexportal.org/home/samplesPage).

## Supplementary Materials

**Materials and Methods**

**Figures S1 – S8**

**Tables S1 – S6**

