## Supplementary material for "Cell type specific genetic regulation of gene expression across human tissues": Kim-Hellmuth et al supplement

### Material and Methods

GTEx data
The GTEx V8 data contains a total of 17,382 RNA-seq samples from 948 post-mortem donors, with 838 donors having genotype data from whole genome sequencing available in a phased analysis freeze VCF(*1*). QTL analyses were based on tissues with at least 70 RNA-seq samples from genotypes donors, corresponding to a total of 15,201 samples, and subsets of these data were used as detailed in the following sections.

xCell cell type enrichment in GTEx
Cell type enrichment scores were computed by running xCell (*2*) on the full TPM gene expression matrix (from RNA-SeQC) of 17,382 RNA-seq samples from the GTEx V8 release, using the `xCellAnalysis` function from the R package.

Benchmarking of xCell cell type estimates
For neutrophils and neurons we compared xCell enrichment scores with estimates from CIBERSORT (*3*), using the built-in LM22 signature matrix or a custom signature matrix (*4*), respectively. For adipocytes, myocytes and keratinocytes, we compared xCell enrichment scores with estimates from the Gene Expression Deconvolution Interactive Tool (GEDIT) (*5*), using default settings and reference data from the Human Body Atlas (*6*) and Swindell et al. (*7*), which are both available on the GEDIT website (http://webtools.mcdb.ucla.edu). For epithelial cells, we compared xCell enrichment scores with estimates from Breschi et al. (*8*). In brief, we employed constrained linear models (lsqlincon function from the pracma R package) to perform cell type deconvolution from gene expression of 368 transverse colon samples. For hepatocytes we were not able to find suitable reference data to compare our xCell estimates against, but tested for correlation with marker genes instead (below). Estimated cell type abundances were compared between methods using Spearman correlation.

For *in silico* simulations, we prepared synthetic cell type mixtures for colon, liver, adipose, skin, muscle, brain and blood tissues. The gene expression of cell types from colon, liver, adipose, skin, muscle was obtained from cell lines from the Human Protein Atlas (*9*), brain cell types were obtained from Zhang et al. (*4*) and blood cell types from Monaco et al. (*10*). xCell was then used to estimate the enrichment of these cell types in the *in silico* mixtures. We varied the proportion of cell types in each of the mixtures using linear combinations of gene expression profiles of each of the cell types. In total, we generated 100 mixtures for each of the tissues and compared the simulated ground truth to xCell enrichment scores using Spearman correlation.

To further assess how well the xCell enrichment scores capture the proportion of specific cell types, we computed the Spearman correlation between the xCell scores and a list of cell type specific marker genes curated from the literature (figs. S1C and S1D). The following markers were used for each cell type: Adipocytes: FASN (*11*); Epithelial cells: CDH1, CLDN7(*12*, *13*); Hepatocytes: AFP(*14*); Keratinocytes: KRT10 (*15*); Myocytes: MYH7, TNNI1 (*16*); Neurons: GAD1 (*17*); Neutrophils: STX3 (*18*).

Histology images were obtained from the GTEx Portal (https://gtexportal.org), and first visually inspected for features matching estimated cell type abundances (e.g., presence of mucosal or muscular layers in transverse colon). The xCell scores also captured differences in histology, as shown for the samples corresponding to the 5th and 95th percentiles of xCell scores in fig. S1C. Next, xCell enrichment scores of 43 cell type-tissue combinations were tested for association with 37 histological phenotypes that were available in the sample attributes for those tissues. The histological phenotypes were harmonized using a test processing methodology described in Breschi et al. (*8*). Briefly, pathology annotations that contain a specific phenotype were identified based on Levenshtein distance. Next, these annotations were processed using part-of-speech tagging with the TextBlob NLP library (https://textblob.readthedocs.io/en/dev/) in order to find dependencies between the words of each annotation and retain those corresponding to the histological phenotypes. Although we were able to recover many annotations with this approach, a small fraction had to be manually curated due part-of-speech mislabeling. Phenotypes needed to have at least three annotated cases in the tissue of interest to be tested. Differences in cell type enrichment score between cases and controls were assessed using the two-sided Wilcoxon Rank-sum test.

<u>Identification of cell type interacting eQTLs and sQTLs</u>

Cell type interaction QTLs were mapped using used a linear regression model with an interaction term accounting for interactions between genotype and cell type enrichment:

$$\mathrm{p} \sim \mathrm{g} + \mathrm{i} + \mathrm{g} \circ \mathrm{i} + \mathrm{C}$$

where $\mathrm{p}$ is the phenotype vector (e.g., gene expression or intron excision ratio), $\mathrm{g}$ is the genotype vector, $\mathrm{i}$ is the inverse normal transformed xCell enrichment score, and the interaction term $\mathrm{g} \circ \mathrm{i}$ corresponds to point-wise multiplication of genotypes and cell type enrichment scores. $\mathrm{C}$ is a matrix of covariates that were also used in regular QTL mapping. These covariates include genotype principal components to correct for population structure and PEER factors. Of note, the interaction term itself is robust to PEERs being correlated with the cell type enrichment scores. Interaction QTLs were identified by testing for the significance of the interaction term, and mapping was performed using tensorQTL (*19*), which computes regression coefficients and p-values for all terms in the model, enabling comparisons of interaction and main effects. Variants within ±1Mb of the TSS of each gene were tested, as for regular QTL mapping. To avoid potential regression outlier effects, we restricted ieQTL mapping to variants with MAF ≥ 0.05 in the samples belonging to each of the top and bottom halves of the enrichment score distribution, for each tissue-cell type combination (using the --maf_threshold_interaction 0.05 option in tensorQTL). For isQTL mapping, this threshold was set to MAF ≥ 0.1. The same filtered and normalized gene expression and splicing phenotype matrices used for regular QTL mapping were used for interaction QTL mapping. To identify genes with at least one significant ieQTL or isQTL (ieGenes or isGenes, respectively), the top nominal p-values for each gene or phenotype was corrected for multiple testing at the gene level using eigenMT (*20*). Significance across genes was computed by adjusting the eigenMT-corrected p-values using Benjamini-Hochberg, and applying a 0.05 FDR threshold. For isQTLs, the p-value corresponding to the top splicing phenotype was selected for each gene-variant pair, and corrected by the number of phenotypes tested ($\tilde{p} = \min{(np, 1)}$), where $\mathrm{n}$ is the number of splicing phenotypes for the gene) prior to running eigenMT. QTL mapping and FDR correction were performed using expression and splicing phenotypes for all biotypes in the GENCODE v26 annotation, but downstream analyses are based on protein coding and lincRNA genes only.

Cell type ieQTL validation using aFC of allele-specific expression data

We used allele-specific expression (ASE) data of eQTL heterozygotes to correlate individual-level allelic fold-change (aFC) of an eQTL with individual-level cell type enrichments. For this analysis, we used the phASER haplotype-based ASE data (with phase preserved across genes) for genes with ≥10 ieQTL heterozygous individuals with ≥8 reads of ASE data per gene and nominally significant ASE aFC ($P < 0.05$). The exact number of ieQTLs tested for ASE aFC validation after these filtering steps is indicated as bar labels in Fig. 2B. Since the number of ieQTLs in many tissue-cell type combinations was too low to compute the π1 statistic (*21*), we used Spearman correlation p-values to assess how many cell type ieQTL show evidence of validation. We report the proportion of ieQTLs with a significant aFC-cell type correlation ($P < 0.05$) for all tested cell type-tissue combinations (Fig. 2B). For 13 cell type-tissue pairs with > 20 ieQTLs at 5% FDR we also calculated the corresponding π1 statistic on the correlation p-values using a fixed lambda of 0.5 (Fig. S5B).

Replication in external data sets

To assess replication of cell type ieQTLs and isQTLs, we examined p-values for matched variant-gene pairs in external cohorts, where we performed the same xCell cell type enrichment analyses and ieQTL and isQTL mapping as described above. Adipocyte and Keratinocyte ieQTLs were tested in TwinsUK adipose and skin tissues respectively (*22*). Neutrophil ieQTLs were tested in whole blood from the GAIT2 study (*23*) and from purified neutrophils (*24*). Neuron ieQTLs and isQTLs were tested in temporal cortex from the Mayo RNA-sequencing study (*25*).

Multi-tissue ieQTL analysis

MASH (*26*) was used to estimate ieQTL activity across 16 epithelial tissues and 13 brain tissues, respectively. We assessed the significance of the top ieQTL-SNP per gene (the SNP with the largest univariate |Z|-statistic across tissues) where ieQTL effect size and standard error was available in all tested tissues. MASH was run in the exchangeable Z (EZ) mode, and 250,000 randomly selected SNP*GENE pairs that were tested across all tissues were used to fit the mash model. Effect size estimates and local false sign rate (LFSR) outputted by MASH were used for QTL magnitude and activity respectively. For pairwise tissue-sharing analysis of ieQTL we considered only iQTLs that were significant (LFSR < 0.05) in at least one of the two tissues. We defined an ieQTL to be shared if they had the same sign and similar magnitude (effect within a factor of 2 of one another), which is implemented in the MASH function `get_pairwise_sharing`.

Tissue specificity analysis of ieQTLs

A logistic regression model of eQTL tissue activity was built to predict whether a eQTL identified in a given discovery tissue is active in a given replication tissue given a set of predictors derived from genomic annotations, and tissue specific gene expression, and chromatin states (*27*). eQTL activity was defined as MASH LFSR < 0.05 in a replication tissue. Basic QC on the eQTL data used to build the model was performed as follows: expression level > 0 in both discovery and replication tissues, eQTL MAF > 0.005 in both discovery and replication tissues, difference in expression level > quantile(0.005) and < quantile (0.995) to exclude the most extreme cases of expression difference. R v3.5.1 was used with speedglm v0.3-2. When plotting model predictor coefficients they were standardized using the standardize R package v0.2.1 so that they could be plotted on the same scale. When reporting AUCs for the model including different sets of features it was trained on eQTLs spanning

chromosomes 1-20 and tested on eQTLS from chromosomes 21 and 22. Otherwise, tissue level AUCs were generated by holding out holding out individual tissues and predicting the activity of eQTLs found in the other 21 tissues in the held-out tissue. In total, 22 tissues were used for the analyses, which were chosen based on having appropriately paired epigenomic state predictions from the ROADMAP Epigenomics Project (*27*). In cases where there were two highly similar tissues (defined by pairwise tissue gene expression clustering), the tissue with the higher sample size was used. Chromatin state sharing was defined as either shared on not shared based on if the ROADMAP Chromatin state prediction was the same (shared) or different (not shared) between the pairwise tissues. The ROADMAP core 15-state model was used (https://egg2.wustl.edu/roadmap/web_portal/chr_state_learning.html).
To test cell type ieQTLs as predictors of tissue specificity we only included cell types with > 20 ieQTLs (FDR > 0.05) in at least one of the tested tissues. Neurons and Hepatocytes had max. 14 and 2 ieQTLs (FDR > 0.05) respectively and were excluded from the analysis. The following predictors were also used in the model: distance between variant and TSS, variant MAF in GTEx, effect size in discovery tissue (aFC), global gene expression correlation between discovery and replication tissue, variant effect prediction, linsight conservation score (*28*), variant is INDEL, Roadmap state and sharing between discovery and replication tissue, variant overlaps Ensembl Regulatory Build TF binding site (in any Ensembl tissue), variant overlaps Ensembl Regulatory Build CTCF binding site (in any Ensembl tissue), variant overlaps Ensembl Regulatory Build DHS site (in any Ensembl tissue), variant overlaps Ensembl Regulatory Build predicted motif site. The Ensembl Regulatory Build annotations were from Zerbino et al. (*29*).

<u>GWAS enrichment analysis</u>
To test whether ieQTLs or isQTLs were enriched for complex disease or trait associations (genome-wide and subthreshold) GWAS hits, we applied an updated version of QTLEnrich (available at https://github.com/segrelabgenomics/QTLEnrich). QTLEnrich assesses the enrichment of top ranked trait associations (GWAS p-value<0.05 used here) amongst a set of significant QTL-variants in a given tissue, accounting for potential confounding factors such as allele frequency, distance to the transcription start site and local level of LD (number of LD proxy variants; $r^2 \geq 0.5$). Briefly, an enrichment p-value was computed for each GWAS-tissue pair tested, as the fraction of 100,000 randomly sampled sets of null variants (of equal size to that of the QTL-variant set). Fold-enrichment was computed as the number of QTL-variants or null variants with a GWAS p-value below 0.05 divided by 5% of the variant set size. An adjusted fold-enrichment was computed for each QTL-variant set, as the fold-enrichment of the QTL-variant set divided by the median fold-enrichment of 1,000 randomly sampled sets of confounder-matched null variants. QTLEnrich was applied to 87 GWAS and GTEx tissues with at least 100 ieQTLs or isQTLs at 40% FDR, testing 23 and 7 tissues respectively. GWAS enrichments among iQTLs at 40% and 10% FDR were comparable (Fig. S8). QTLEnrich was also applied to 87 GWAS and 49 GTEx tissues for QTLs at 5% FDR. To compare GWAS enrichment among iQTLs vs standard QTLs, we matched tissue-trait pairs for standard QTLs at 5% FDR. The most significant QTL variant per ieGene, isGene, eGene, or sGene was used to reduce potential inflation of enrichment due to local LD. Bonferroni correction was used to determine significant trait-tissue pairs, and the adjusted fold-enrichment was used as the test statistic to rank significant tissues based on their enrichment, as it corrects for enrichment of trait associations amongst matched null variants.

<u>Cell type iQTL-GWAS colocalization analysis</u>

Colocalization analysis was conducted using the coloc R package (*30*). Coloc uses summary statistics from QTL and GWAS studies in a Bayesian framework to identify GWAS signals that colocalize with QTLs. To maximize our discovery power, we ran coloc for all cell type ieGenes at FDR < 0.4 and 87 GWAS traits (table S3). All variants of the *cis*-QTL region (+/- 1 MB of the TSS of an ieGene) that were available for both the QTL and the GWAS trait were used in the function `coloc.abf()` with either *cis*-ieQTL or corresponding cis-eQTL p-values and GWAS effect size estimates and their variances. Given the high sensitivity of colocalization results to the choice of priors, we use model-based priors computed with enloc (*31*). A complete list of priors for each of the 49 tissues and 87 GWAS trait combinations is provided in table S6 for *cis*-eQTLs and *cis*-sQTLs. The same model-based priors used for *cis*-eQTLs were used for *cis*-ieQTLs assuming that regular *cis*-eQTLs reflect the average signal of all ieQTLs for a particular gene. The corresponding prior can thus be interpreted as the average prior of all ieQTLs for that gene and can be used as an approximated prior for each individual cell type ieQTL. We defined an ieGene or eGene as having evidence of colocalization when posterior probability of colocalization (PP4) was higher than 0.5. All coloc results with PP4 ≥ 0.50 are reported in table S5.

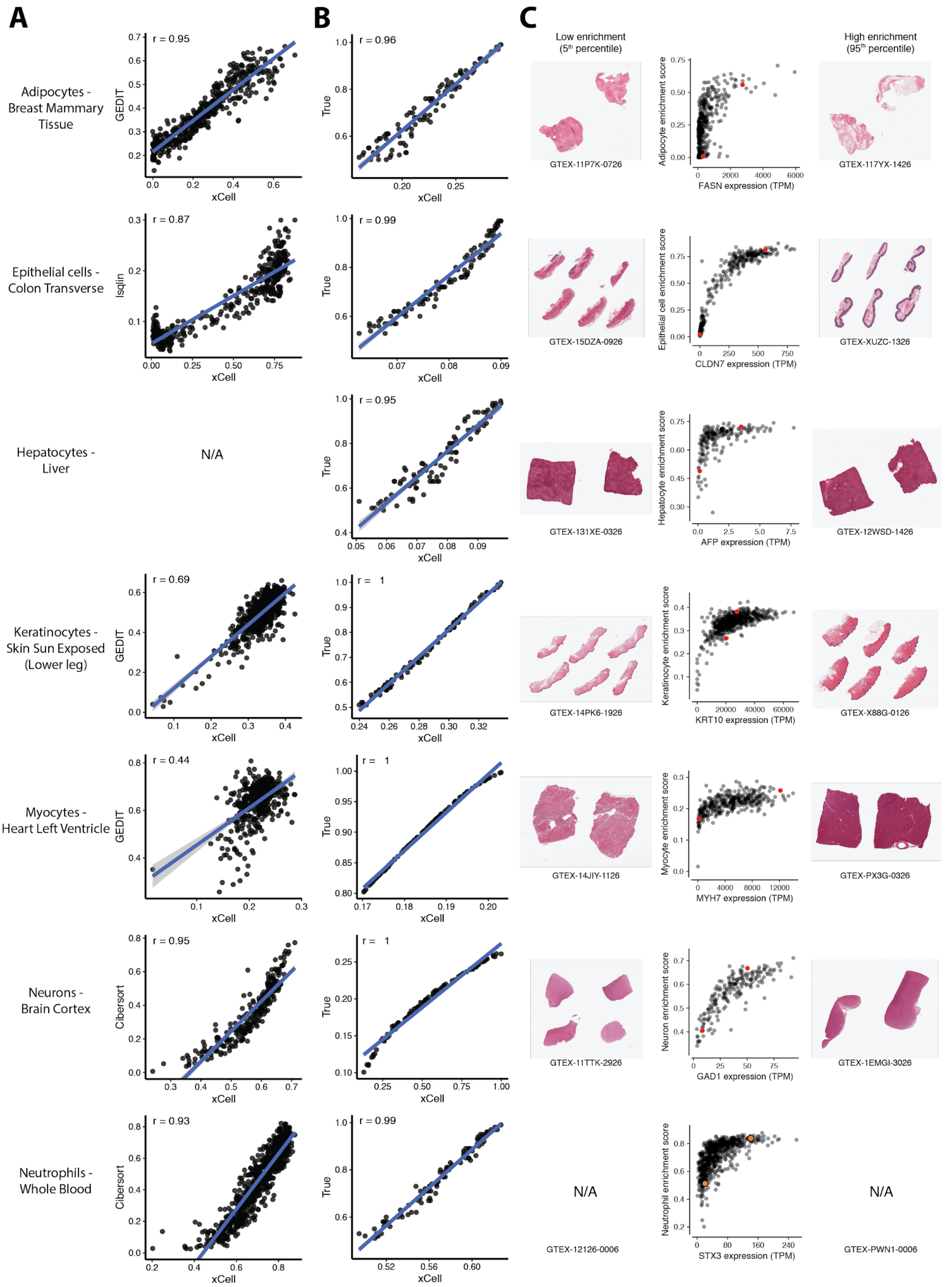
A
B
C
Adipocytes - Breast Mammary Tissue
r = 0.95
GEDIT
xCell
r = 0.96
True
Epithelial cells - Colon Transverse
r = 0.87
lsqlin
r = 0.99
Hepatocytes - Liver
N/A
r = 0.95
Keratinocytes - Skin Sun Exposed (Lower leg)
r = 0.69
r = 1
Myocytes - Heart Left Ventricle
r = 0.44
Neurons - Brain Cortex
r = 0.95
Cibersort
Neutrophils - Whole Blood
r = 0.93
r = 0.99
Low enrichment (5th percentile)
High enrichment (95th percentile)
GTEX-11P7K-0726
Adipocyte enrichment score
FASN expression (TPM)
GTEX-117YX-1426
GTEX-15DZA-0926
Epithelial cell enrichment score
CLDN7 expression (TPM)
GTEX-XUZC-1326
GTEX-131XE-0326
Hepatocyte enrichment score
AFP expression (TPM)
GTEX-12WSD-1426
GTEX-14PK6-1926
Keratinocyte enrichment score
KRT10 expression (TPM)
GTEX-X88G-0126
GTEX-14JIY-1126
Myocyte enrichment score
MYH7 expression (TPM)
GTEX-PX3G-0326
GTEX-11TTK-2926
Neuron enrichment score
GAD1 expression (TPM)
GTEX-1EMGI-3026
N/A
GTEX-12126-0006
Neutrophil enrichment score
STX3 expression (TPM)
N/A
GTEX-PWN1-0006

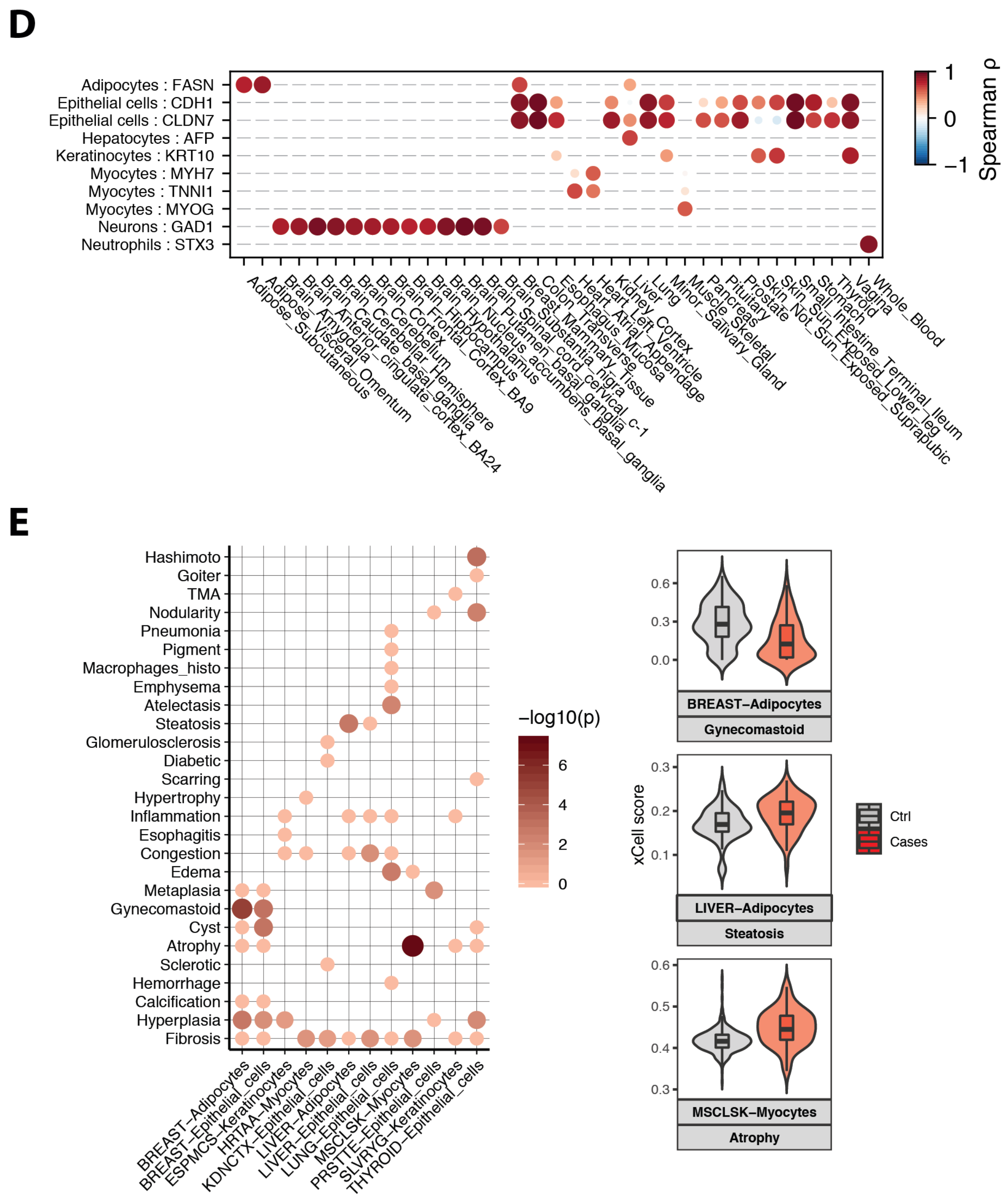


**Fig. S1. Benchmarking of xCell cell type enrichment estimates.** (**A**) Concordance rates between xCell and other methods. Neuron and neutrophil estimates were generated using Cibersort; adipocyte, keratinocyte and myocyte estimates were generated using GEDIT and epithelial cell estimates were generated using a linear model (see Methods). The tissue where each cell type was estimated is indicated above each plot. Sample sizes for each tissue are shown in the top left corner [will be added] and Spearman correlation coefficients are shown in the top right corner. (**B**) in silico simulation using synthetic cell type mixtures for colon, liver, adipose, skin, muscle, brain and blood tissues. (**C**) Histology images corresponding to samples with xCell estimates in the 5$^{th}$ and 95$^{th}$ percentile, and correlation between xCell estimates and cell type marker genes. The red dots indicate the 5$^{th}$ and 95$^{th}$ percentile samples. (**D**) Spearman correlation between expression of cell type-specific marker genes and xCell enrichment scores for combinations of cell types and tissues used in this paper (other, non-tested combinations are indicated by gray lines). Normalized expression data for mapping QTLs was used (see Methods). (**E**) Cell type estimates of 42 cell type-tissue pairs tested for association with 37 histological phenotypes that were available for those tissues (Methods). Phenotypes needed to have at least 3 annotated cases in the tissue of interest to be tested. In rows, phenotypes that were significantly associated in at least one cell type-tissue pair are shown. In columns, cell type-pairs that were significantly associated with at least one phenotype are shown. The color and size of each dot indicate Wilcoxon Rank-sum p-values. Three examples are shown in the right-hand side panel. Adipocytes in Breast tissue are depleted in gynecomastia samples compared to controls. Adipocytes in liver cases with steatosis are elevated compared to control. Myocytes in atrophic skeletal muscle are elevated.

A

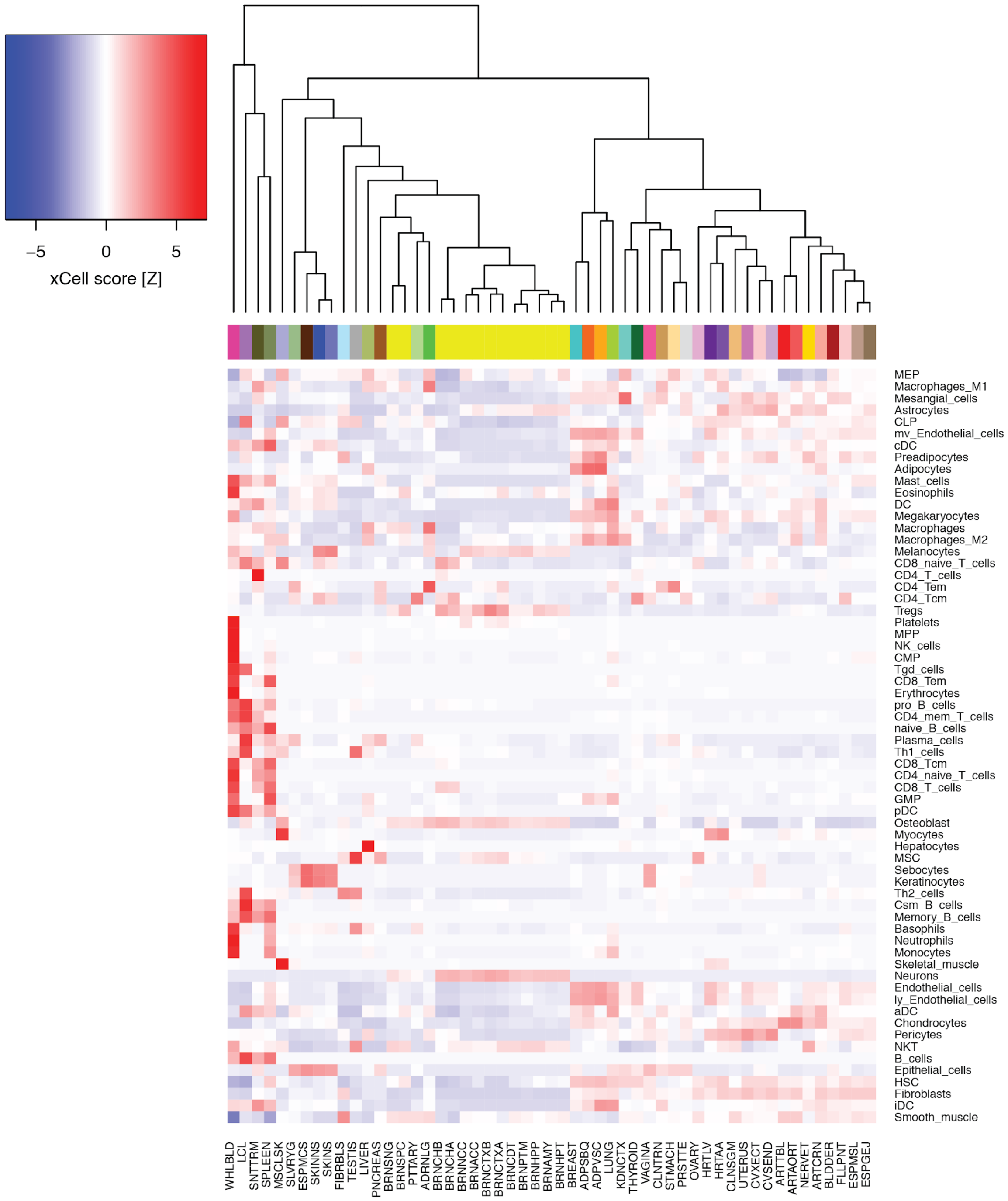

−5
0
5
xCell score [Z]
MEP
Macrophages_M1
Mesangial_cells
Astrocytes
CLP
mv_Endothelial_cells
cDC
Preadipocytes
Adipocytes
Mast_cells
Eosinophils
DC
Megakaryocytes
Macrophages
Macrophages_M2
Melanocytes
CD8_naive_T_cells
CD4_T_cells
CD4_Tem
CD4_Tcm
Tregs
Platelets
MPP
NK_cells
CMP
Tgd_cells
CD8_Tem
Erythrocytes
pro_B_cells
CD4_mem_T_cells
naive_B_cells
Plasma_cells
Th1_cells
CD8_Tcm
CD4_naive_T_cells
CD8_T_cells
GMP
pDC
Osteoblast
Myocytes
Hepatocytes
MSC
Sebocytes
Keratinocytes
Th2_cells
Csm_B_cells
Memory_B_cells
Basophils
Neutrophils
Monocytes
Skeletal_muscle
Neurons
Endothelial_cells
ly_Endothelial_cells
aDC
Chondrocytes
Pericytes
NKT
B_cells
Epithelial_cells
HSC
Fibroblasts
iDC
Smooth_muscle
WHLBLD
LCL
SNTTRM
SPLEEN
MSCLSK
SLVRYG
ESPMCS
SKINNS
SKINS
FIBRBLS
TESTIS
LIVER
PNCREAS
BRNSNG
BRNSPC
PTTARY
ADRNLG
BRNCHB
BRNCHA
BRNNCC
BRNACC
BRNCTXB
BRNCTXA
BRNCDT
BRNPTM
BRNHPP
BRNAMY
BRNHPT
BREAST
ADPSBQ
ADPVSC
LUNG
KDNCTX
THYROID
VAGINA
CLNTRN
STMACH
PRSTTE
OVARY
HRTLV
HRTAA
CLNSGM
UTERUS
CVXECT
CVSEND
ARTTBL
ARTAORT
NERVET
ARTCRN
BLDDER
FLLPNT
ESPMSL
ESPGEJ

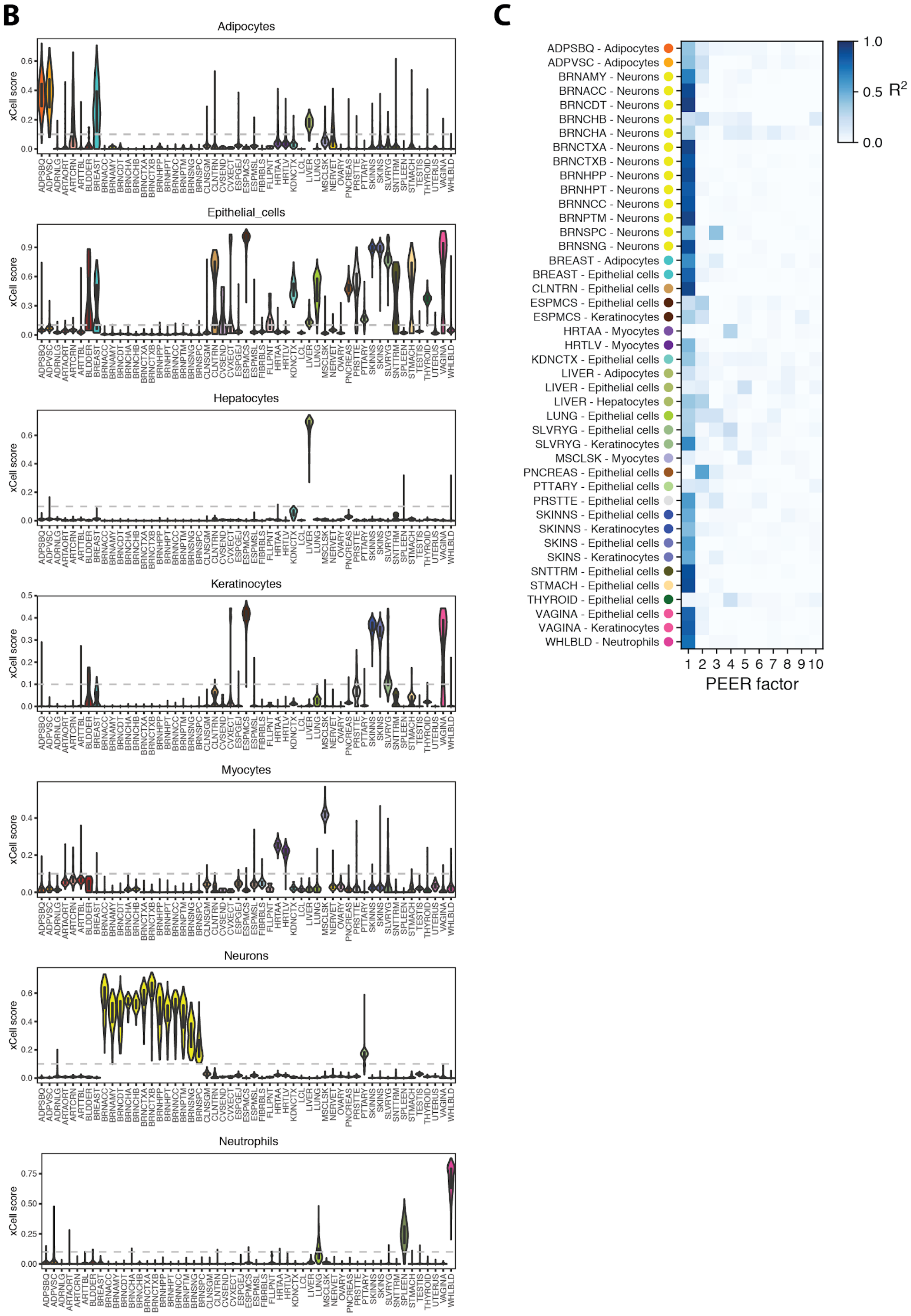


**Fig. S2. xCell enrichment scores across 49 tissues. (A)** Mean xCell enrichment scores across tissues. **(B)** xCell estimates of seven cell types across 49 tissues. For each cell type, interaction eQTL analysis was performed only in tissues where the cell type had a median xCell score > 0.1 (dashed horizontal line). (**C**) Correlation between xCell enrichment scores for the cell types indicated on the y-axis and the first ten PEER factors computed for each tissue, showing that the top PEER factors capture cell type heterogeneity.

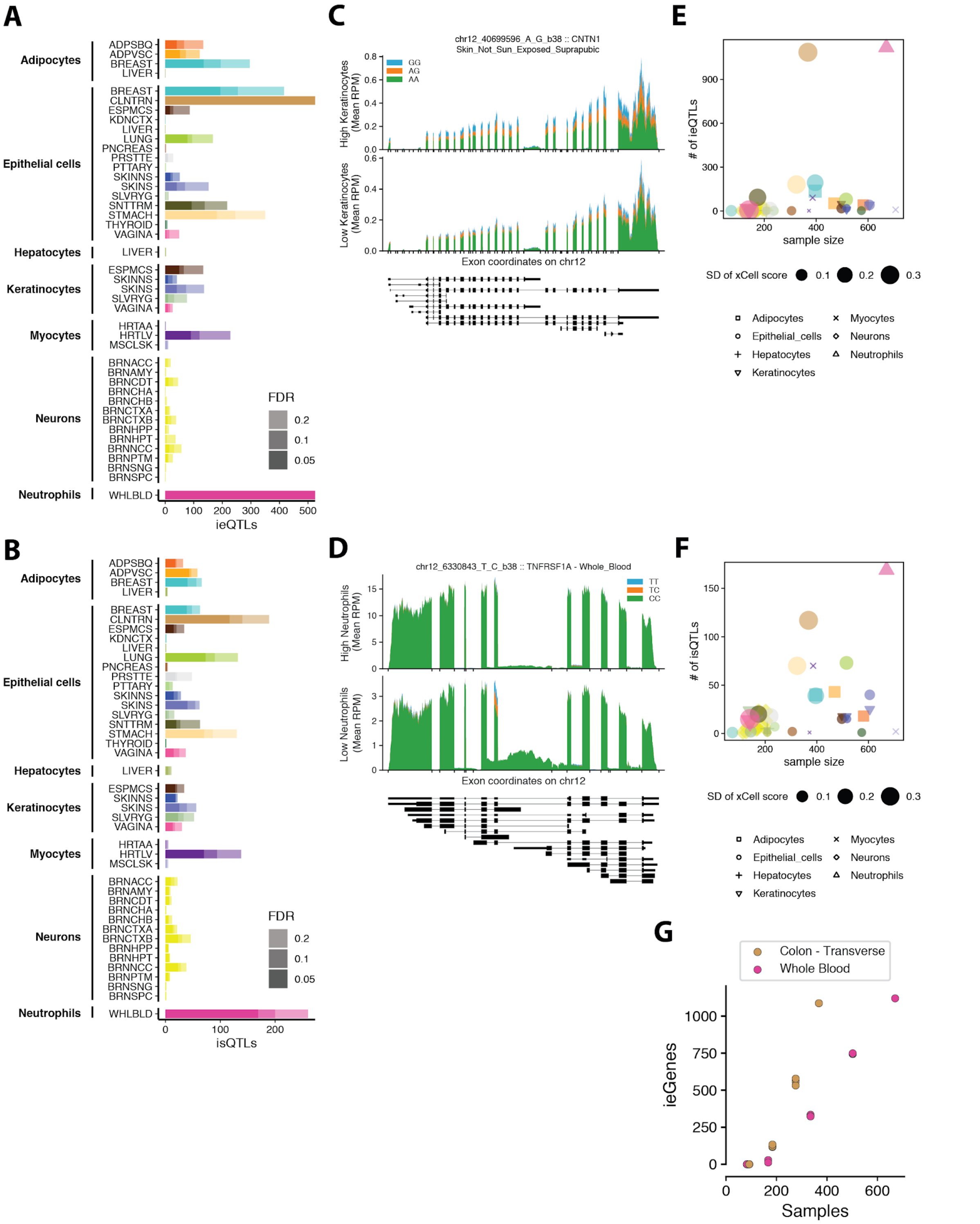


**Fig. S3. Cell type ieQTL discovery.** (**A**) Number of cell type ieQTL discovered in each cell type-tissue pair at indicated FDR thresholds. x-axis is truncated at 500. The numbers of neutrophil-ieQTLs in Whole Blood were 1120/ 1380/ 1781 (FDR 0.05/ 0.1/ 0.2). The numbers of epithelial cell-ieQTLs in Colon Transverse were 1087/ 1513/ 2169 (FDR 0.05/ 0.1/ 0.2). (**B**) Number of cell type isQTLs discovered in each cell type-tissue pair at indicated FDR thresholds. (**C**) ieQTL pileup for CNTN1 in not sun-exposed skin. (**D**) isQTL pileup for TNFRSF1A in whole blood. (**E**) Number of cell type ieQTLs per tissue, as a function of sample size and variance of cell type estimates. The size reflects the standard deviation of cell type estimates in the corresponding tissue. Cell types are depicted as symbols. (**F**) The number of cell type isQTLs per tissue, as a function of sample size and variance of cell type estimates. (**C**) Downsampling analysis of cell type ieQTLs in whole blood and transverse colon.

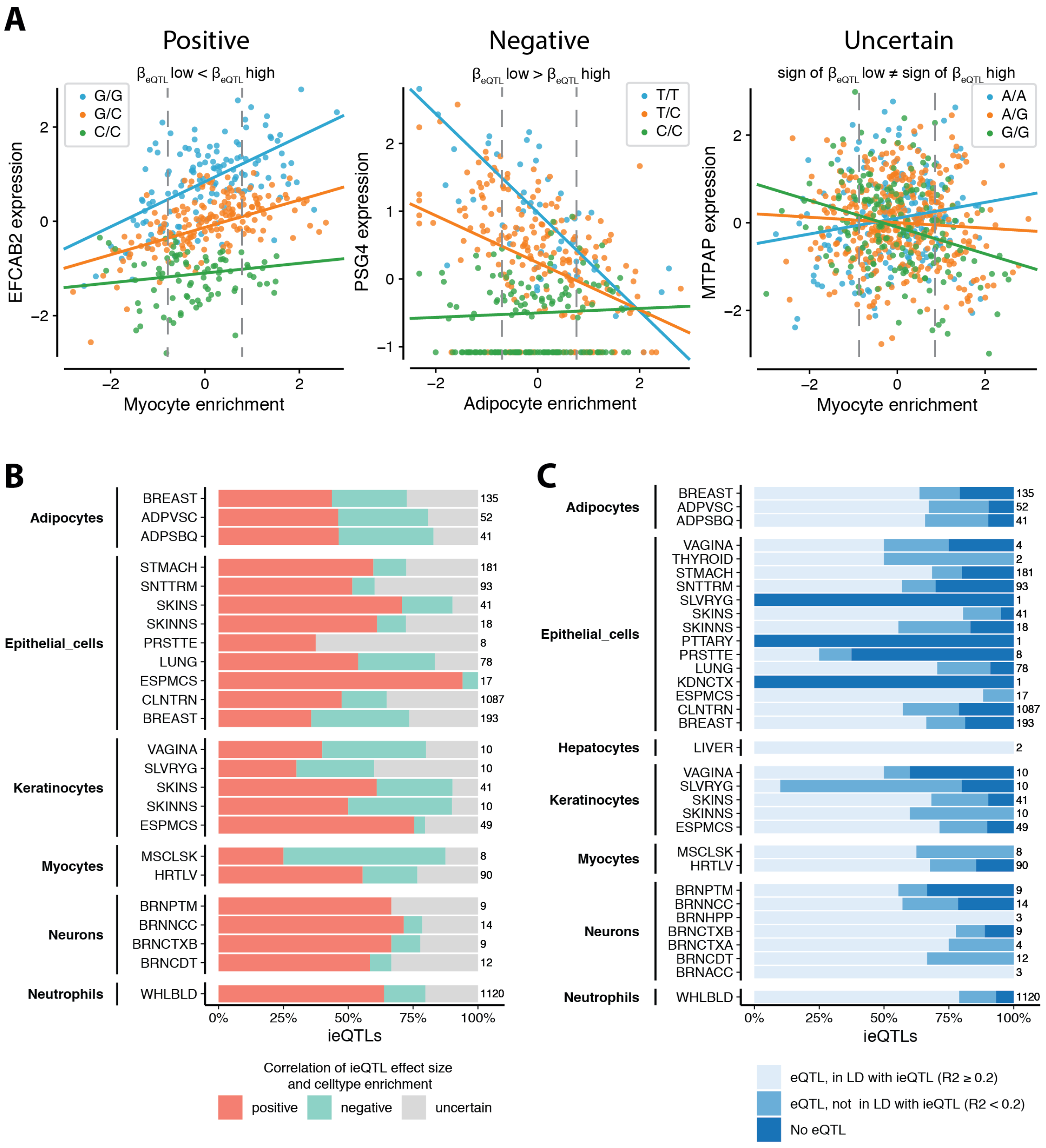


**Fig. S4. Correlation of ieQTLs and cell type estimates.** (**A**) ieQTL examples that are either positively (left) or negatively (middle) correlated with cell type estimates, or where cell type correlation is uncertain (right). To categorize ieQTLs into these groups genotype main effects at low (25$^{th}$ percentile) vs high (75$^{th}$ percentile) cell type enrichment were compared. ieQTLs with "positive" cell type correlation show an increase of the genotype main effect from low to high cell enrichments. ieQTLs with "negative" cell type correlation show a decrease and the "uncertain" group contains ieQTLs where the sign flips between low and high cell type enrichments. (**B**) Stacked bar plots of proportion of cell type ieQTLs (min. 5 ieQTLs with FDR < 0.05) that show positive, negative or uncertain cell type correlation. Numbers at the end of each stacked bar plot indicate the total number of cell type ieQTLs per tissue at FDR 0.05. (**C**) Proportion of cell type ieQTLs discovered in 33 tissue-cell type pairs (min. 1 ieQTL with FDR < 0.05), with shading indicating whether the ieQTL was discovered by *cis*-eQTL analysis in bulk tissue.

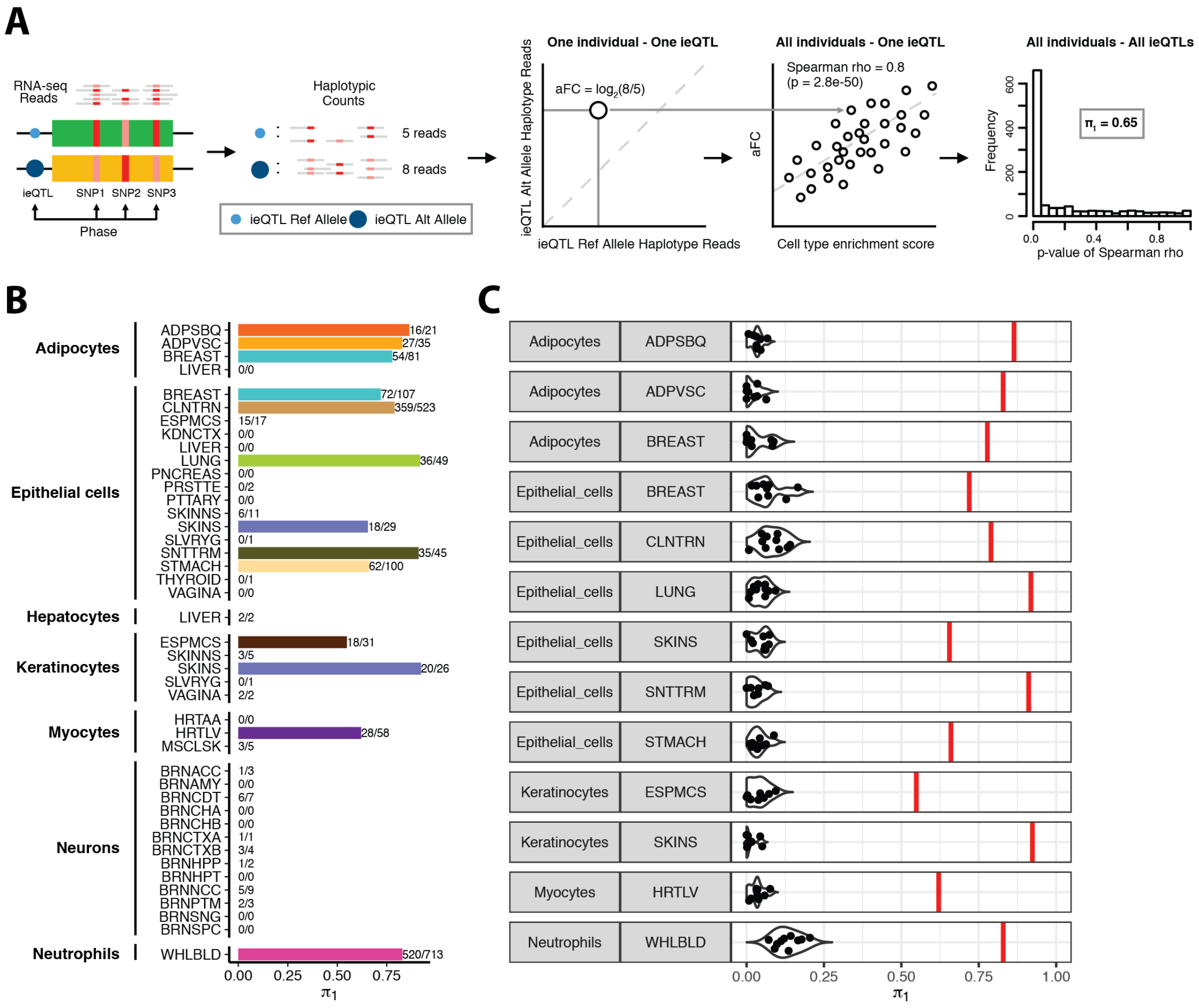


**Fig. S5. Validation of cell type ieQTLs using correlation of ASE aFC and cell type abundance estimates.** (**A**) Schematic representation of the validation approach. Using haplotype-based ASE data read counts can be summed up for each haplotype of an individual who is heterozygous for the ieQTL-SNP. The effect size or aFC is calculated for each heterozygous individual. For each ieQTL correlation of aFC and cell type enrichment scores can be used to assess if the cell type-dependent increase or decrease of the eQTL effect is also seen in ASE aFC data. π1 statistic can be used to report overall validation rate. (**B**) π1 validation rate for cell type-tissue pairs with > 20 ieQTLs. Bar labels indicate the number of ieQTLs with validation/number of ieQTLs tested. (**C**) π1 null distributions for 10 sets of 1000 randomly sampled ieQTLs for each of the 13 cell type-tissue combinations shown in (B).

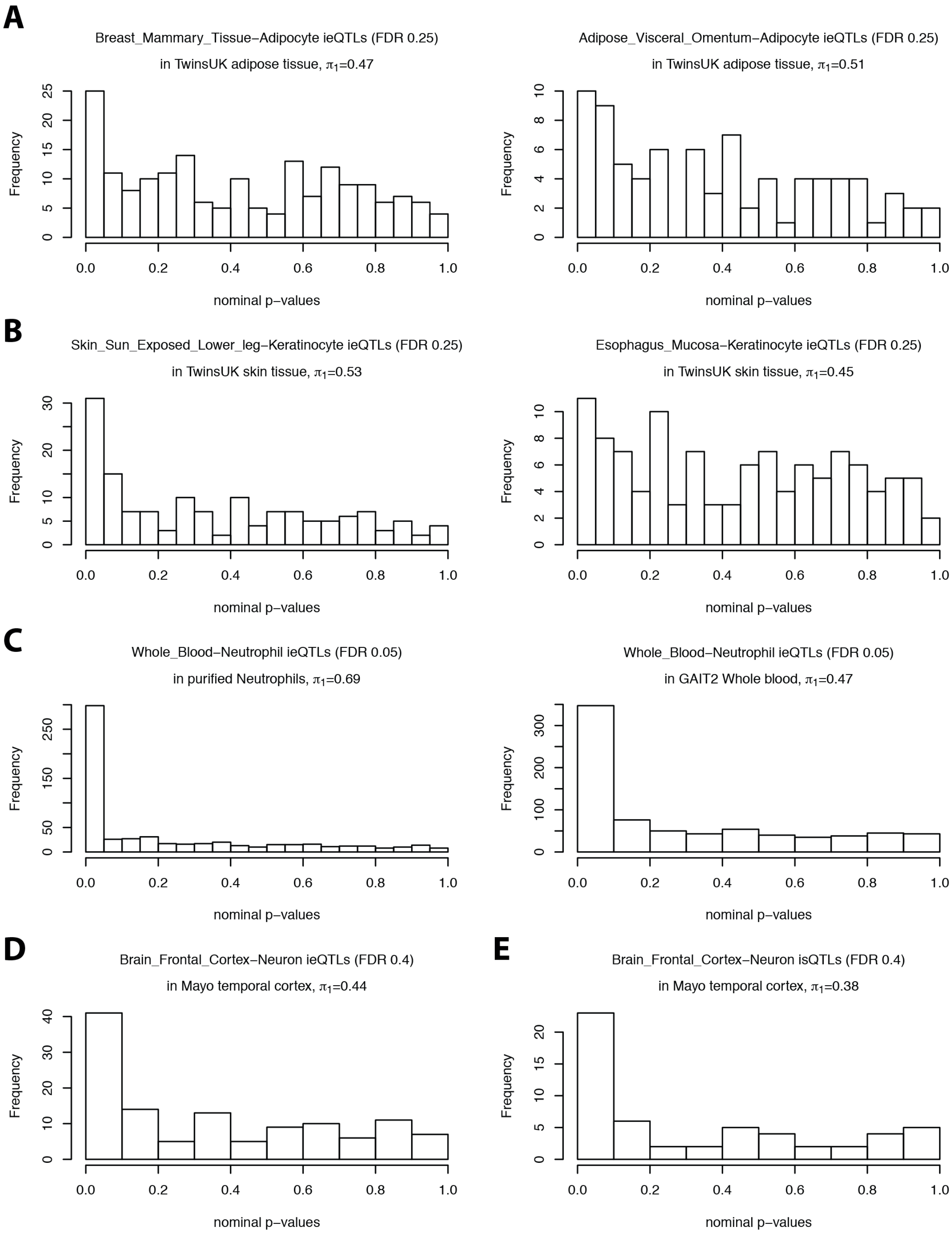


**Fig. S6. Replication of ieQTLs and isQTLs in external studies.** We examined p-values for matched variant-gene pairs in external cohorts, where we performed xCell cell type enrichment analysis followed by interaction QTL mapping. To generate robust π1 replication estimates we used different FDR thresholds indicated in the header to test at least 100 ieQTLs. (**A**) Replication of adipocyte ieQTLs from two different GTEx tissues in TwinsUK adipose tissue (*22*). (**B**) Replication of keratinocyte ieQTLs from two different GTEx tissues in TwinsUK skin tissue. (**C**) Replication of neutrophil ieQTLs in purified neutrophils from (*24*) and the GAIT2 study (*23*). Replication of neuron ieQTLs (**D**) and isQTLs (**E**) in the Mayo RNA-seq study (*25*). For neuron isQTLs only 69 isQTL were available at FDR 0.4.

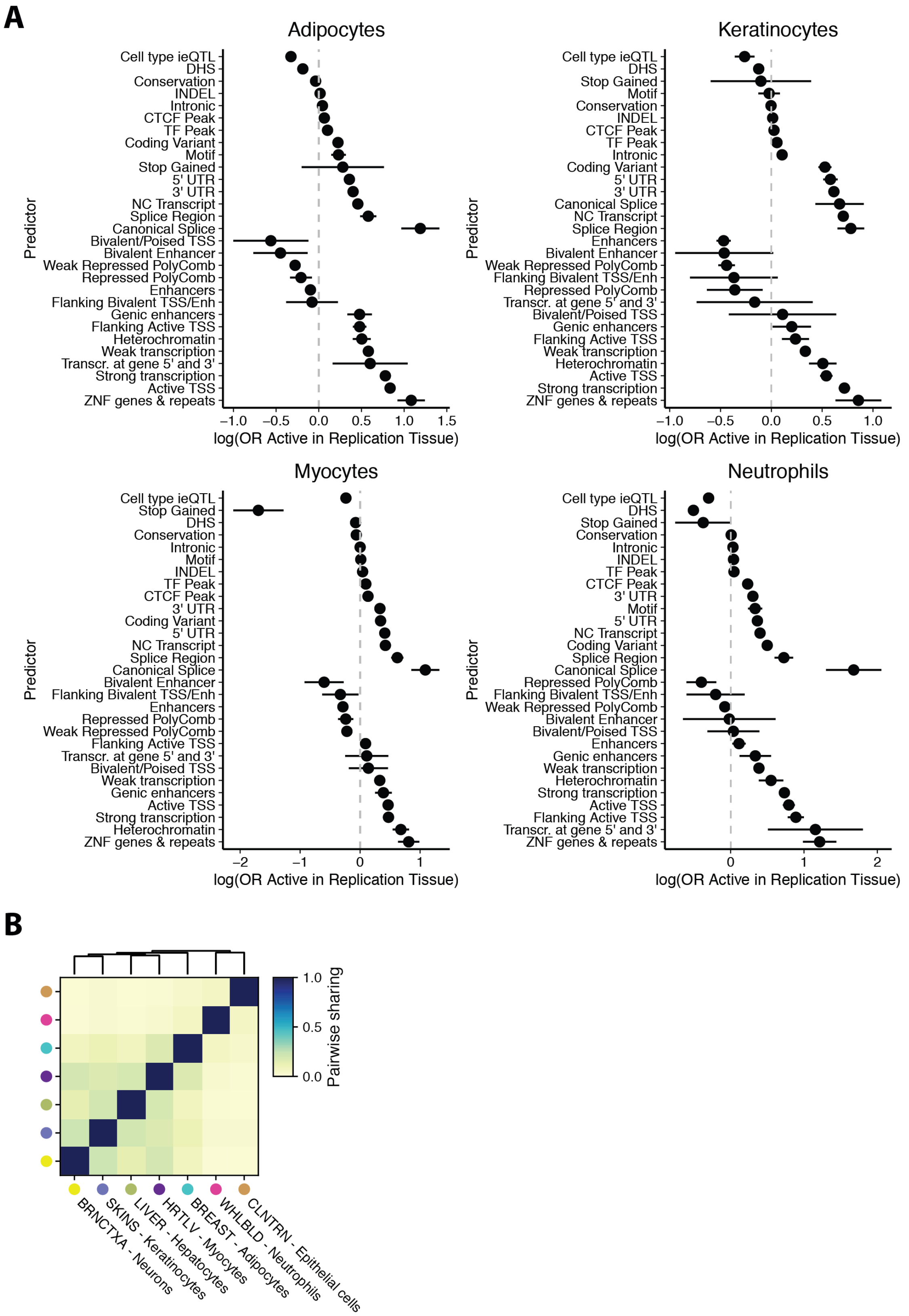


**Fig. S7. Mechanism of eQTL tissue-specificity.** (**A**) Coefficients from logistic regression models of *cis*-eQTL tissue sharing where cell ieQTL status of the four remaining cell types with > 20 ieQTLs is one of the predictors (related to Fig. 3A). All significant top *cis*-eQTLs per tissue were annotated based on if they were also a significant ieQTL for a given cell type. The coefficients represent the log(odds ratio) that an eQTL is active in a replication tissue given a predictor. Bars represent the 95% confidence interval. (**B**) Pairwise sharing by magnitude and sign (from MASH) of ieQTLs across seven different tissue-cell type pairs.

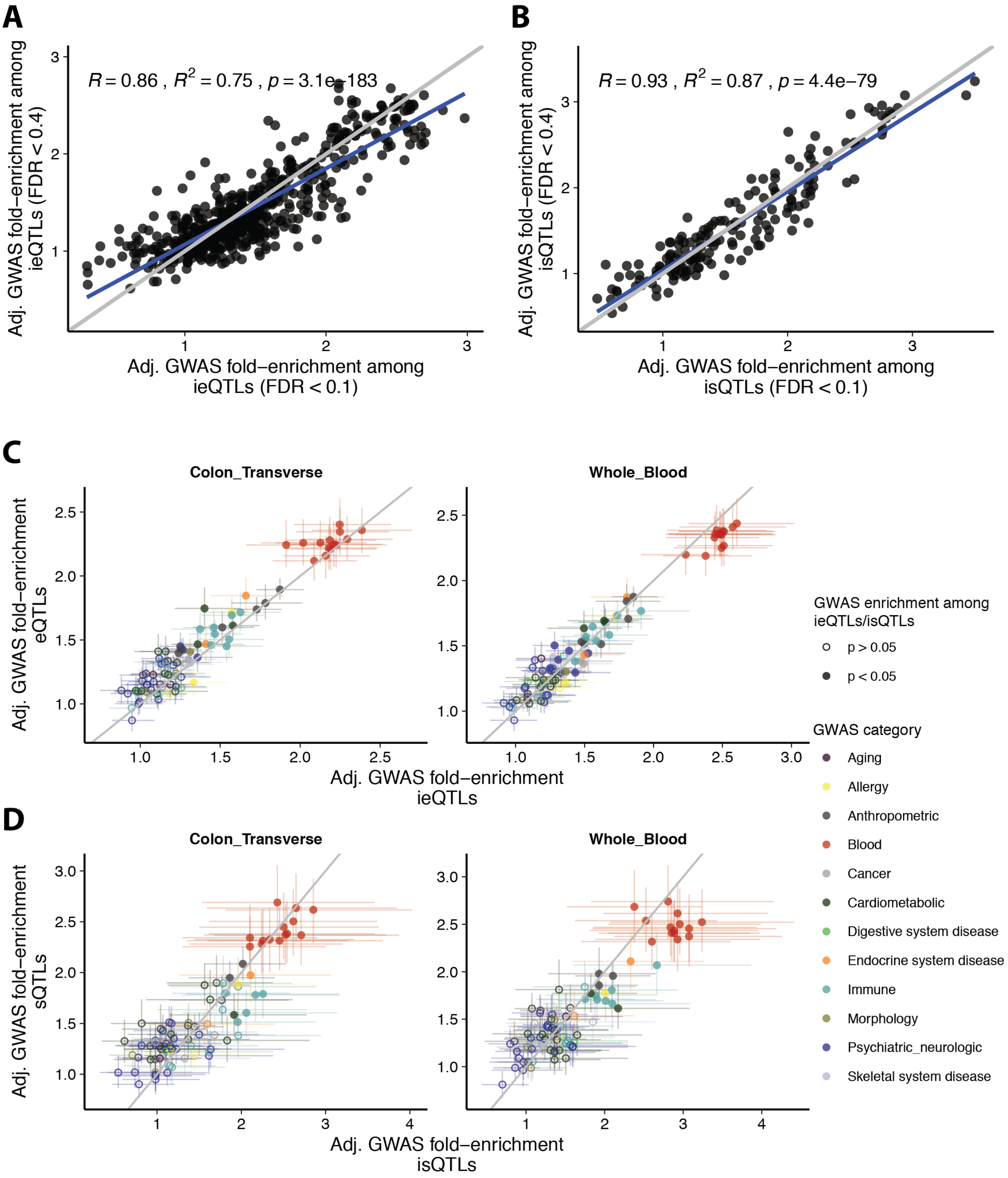


**Fig. S8. QTLEnrich analysis of iQTLs.** Comparison of GWAS adjusted fold-enrichment among ieQTLs (**A**) and isQTLs (**B**) at 40% FDR vs 10% FDR. Blue line represents the fit, grey line indicates the diagonal. (**C**)+(**D**) Same plots as Fig.4B including 95% confidence intervals for ieQTLs (**C**) and isQTLs (**D**) respectively. Adjusted GWAS fold-enrichments of 87 GWAS traits among iQTLs on the x-axis and standard QTLs on the y-axis. Filled circles indicate significant GWAS enrichment among iQTLs at $p < 0.05$ (Bonferroni-corrected). Colors represent GWAS categories of the 87 GWAS traits (see table S3).
